# HPV E6 upregulates MARCHF8 ubiquitin ligase and inhibits apoptosis by degrading the death receptors in head and neck cancer

**DOI:** 10.1101/2022.09.07.507063

**Authors:** Mohamed I. Khalil, Canchai Yang, Lexi Vu, Smriti Chadha, Craig Welbon, Claire D. James, Iain M. Morgan, William C. Spanos, Dohun Pyeon

**Affiliations:** Department of Microbiology and Molecular Genetics, Michigan State University, East Lansing, Michigan, USA; Department of Molecular Biology, National Research Centre, El-Buhouth St., Cairo, Egypt; Cancer Biology and Immunotherapies Group, Sanford Research, Sioux Falls, South Dakota, USA; Philips Institute for Oral Health Research, School of Dentistry, Virginia Commonwealth University, Richmond, Virginia, USA

## Abstract

The membrane-associated RING-CH-type finger ubiquitin ligase MARCHF8 is a human homolog of the viral ubiquitin ligases Kaposi’s sarcoma herpesvirus K3 and K5 that promote host immune evasion. Previous studies have shown that MARCHF8 ubiquitinates several immune receptors, such as the major histocompatibility complex II and CD86. While human papillomavirus (HPV) does not encode any ubiquitin ligase, the viral oncoproteins E6 and E7 are known to regulate host ubiquitin ligases. Here, we report that MARCHF8 expression is upregulated in HPV-positive head and neck cancer patients but not in HPV-negative head and neck cancer patients compared to normal individuals. MARCHF8 expression is highly upregulated by HPV oncoprotein E6-induced MYC/MAX transcriptional activation. The knockdown of MARCHF8 expression in human HPV-positive HNC cells restores cell surface expression of the tumor necrosis factor receptor superfamily (TNFRSF) death receptors, FAS, TRAIL-R1, and TRAIL-R2, and enhances apoptosis. MARCHF8 protein directly interacts with and ubiquitinates the TNFRSF death receptors. Further, MARCHF8 knockout in mouse oral cancer cells expressing HPV16 E6 and E7 augments cancer cell apoptosis and suppresses tumor growth in vivo. Our findings suggest that HPV inhibits host cell apoptosis by upregulating MARCHF8 and degrading TNFRSF death receptors in HPV-positive HNC cells.

**IMPORTANCE:** Since host cell survival is essential for viruses to replicate persistently, many viruses have evolved to prevent host cell apoptosis. The human papillomavirus (HPV) oncoproteins are known to dysregulate proapoptotic proteins. However, our understanding of detailed mechanisms for HPV to inhibit apoptosis is limited. Here, we report that HPV E6 induces transcription of the membrane-associated ubiquitin ligase MARCHF8, which is upregulated in HPV-positive head and neck cancer. MARCHF8 ubiquitinates the tumor necrosis factor receptor superfamily (TNFRSF) death receptors, FAS, TRAIL-R1, and TRAIL-R2 for degradation. We further revealed that downregulation of the death receptors by MARCHF8 prevents cancer cell apoptosis and that knockout of MARCHF8 expression significantly inhibits in vivo tumor growth and enhances tumor-free survival of mice transplanted with mouse oral cancer cells expressing HPV16 E6 and E7.These results suggest that virus-induced degradation of death receptors leads to cancer cell survival in HPV-positive head and neck cancer.

## INTRODUCTION

High-risk human papillomavirus (HPV) is associated with about 5% of all human cancers, including ∼25% of head and neck cancers (HNCs). HPV-positive HNC (HPV+ HNC) arises mainly in the oropharynx, while HPV-negative HNC (HPV-HNC), linked to smoking and drinking, develops in various mouth and throat regions (1, 2). The HPV+ HNC incidence has increased dramatically in recent decades (3–5). The HPV oncoproteins E6 and E7 contribute to fundamental oncogenic mechanisms in HPV+ HNC development that require multiple oncogenic driver mutations in HPV-HNC (6, 7).

Virus-induced ubiquitination and degradation of death receptors are a potent mechanism for host cell survival and viral replication (8). Several DNA viruses downregulate the surface expression of the tumor necrosis factor receptor superfamily (TNFRSF) death receptors such as FAS and tumor necrosis factor (TNF)-related apoptosis-inducing ligand receptors (TRAIL-R1 and TRAIL-R2). For example, adenoviral E3 and human cytomegalovirus UL141 proteins inhibit apoptosis by inducing internalization and lysosomal degradation of the TNFRSF death receptors (9–11). In addition, HPV16 E6 protein binds to the FAS-associated death domain and prevents FAS-induced apoptosis (12).

The membrane-associated RING-CH-type finger (MARCHF) proteins are a subfamily of the RING-type E3 ubiquitin ligase family (13). The MARCHF proteins contain a C4HC3-type RING domain, initially identified in K3 and K5 ubiquitin ligases of Kaposi’s sarcoma-associated herpesvirus (KSHV) (14–16). MARCHF8, a MARCHF family member, is expressed in various tissue types and localizes in the plasma and endosome membranes (17). Similar to KSHV K3 and K5 proteins, MARCHF8 ubiquitinates immunoreceptors such as the major histocompatibility complex II (MHC-II) (18) and CD86 (19), the cell adhesion molecules CD98 and CD44 (20), and the death receptor TRAIL-R1 (a.k.a., death receptor 4 and TNFRSF10A) (21). Previous studies have shown that *MARCHF8* expression is upregulated in esophageal, colorectal, and gastric cancers (22–24). However, the role of MARCHF8 in HPV-associated cancers is largely elusive despite its importance in regulating immune and death receptors.

Here, we report a novel mechanism of HPV oncoprotein E6 for the viral escape of host cell apoptosis through upregulation of *MARCHF8* expression. HPV16 E6 activates MARCHF8 transcription through MYC/MAX activity. Increased MARCHF8 protein in HPV+ HNC cells binds to and ubiquitinates FAS, TRAIL-R1, and TRAIL-R2 proteins. Knockdown and knockout of *MARCHF8* expression in HPV+ HNC cells significantly enhances apoptosis and attenuates in vivo tumor growth. Our findings provide a novel insight into virus-induced evasion of host cell apoptosis and a potential therapeutic target to treat HPV+ HNC.

## RESULTS

### MARCHF8 expression is significantly upregulated in HPV+ HNC by HPV oncoproteins

To determine if *MARCHF8* expression levels are altered in HPV+ and HPV-HNC patients, we analyzed our gene expression data from HPV+ (*n* = 16) and HPV-(*n* = 26) HNC patients and normal individuals (*n* = 12)(25). Our result showed that MARCHF8 mRNA expression in HPV+ HNC patients, but not in HPV-HNC patients, is significantly upregulated compared to the normal individuals (**Fig. 1A**). Although several HPV-HNC patients also showed high MARCHF8 mRNA levels, there were no statistically significant differences compared to normal individuals and HPV+ HNC patients (**Fig. 1A**). We also measured MARCHF8 mRNA levels in HPV+ HNC cell lines (SCC-2, SCC90, and SCC-152), HPV-HNC cell lines (SCC-1, SCC-9, and SCC-19), and normal hTERT immortalized keratinocytes (N/Tert-1) by RT-qPCR. Our results showed significantly higher levels of MARCHF8 mRNA in all HPV+ HNC cells compared to N/Tert-1 and HPV-HNC cells (**Fig. 1B**). Next, to assess if HPV16 E6 and/or E7 are sufficient to upregulate *MARCHF8* expression, we established in N/Tert-1 cells expressing HPV16 E6 and/or E7 using lentiviral transduction and puromycin selection. MARCHF8 mRNA levels were measured in N/Tert-1 vector, N/Tert-1 E6, N/Tert-1 E7, and N/Tert-1 E6E7 cells by RT-qPCR. The results showed that expression of either or both E6 and E7 was sufficient to significantly upregulate MARCHF8 mRNA expression in N/Tert-1 cells (**Fig. 1C**). Western blotting analyses demonstrated that MARCHF8 protein levels are also significantly higher in HPV+ HNC (**Fig. 1D and 1E**) and E6/E7 expressing N/Tert-1 cells (**Fig. 1F and 1G**) compared to N/Tert-1. These results suggest that *MARCHF8* expression is upregulated in HPV+ HNC cells by the HPV oncoproteins E6 and E7.

**FIG. 1.**
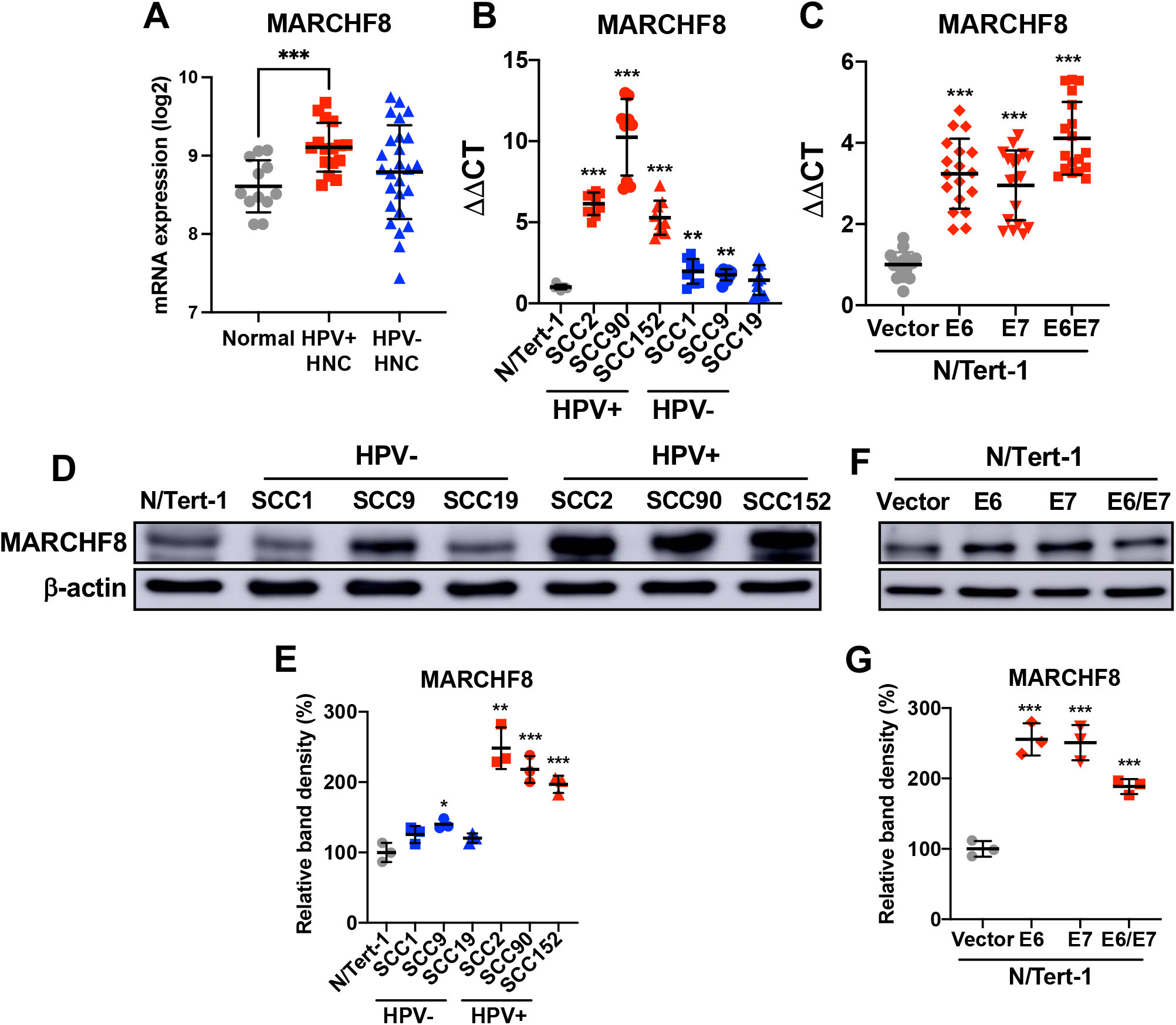
*MARCHF8* expression is upregulated in HPV+ HNC. MARCHF8 mRNA expression levels in microdissected human tissue samples from HPV+ (*n* = 16) and HPV-(*n* = 26) HNC patients, and normal individuals (*n* = 12) were analyzed using our previous gene expression data (GSE6791) (25) and shown as fluorescence intensity (log2) (**A**). The MARCHF8 mRNA expression was validated in normal (N/Tert-1), HPV+ HNC (SCC-2, SCC-90, and SCC-152), and HPV-HNC (SCC-1, SCC-9, and SCC-19) cells (**B**) by RT-qPCR. MARCHF8 mRNA levels were determined in N/Tert-1 cells expressing E6 and/or E7 by RT-qPCR (**C**). The data shown are ΔΔCT values normalized by the GAPDH mRNA level as an internal control. MARCHF8 protein levels were determined in HPV+ and HPV-HNC cells (**D and E**) and N/Tert-1 cells expressing HPV16 E6, E7, or E6 and E7 (**F and G**) by western blotting. β-actin was used as a loading control. The relative band density was quantified using the NIH ImageJ program (**E and G**). All experiments were repeated at least three times, and the data shown are means ± SD. *P* values were determined by Student’s *t*-test. \**p* < 0.05, \*\**p* < 0.01, \*\*\**p* < 0.001.

### The HPV oncoprotein E6 induces *MARCHF8* promoter activity through the MYC/MAX complex

To assess the transcriptional regulation of MARCHF8 mRNA expression by HPV oncoproteins, *MARCHF8* promoter activity was analyzed in HPV+ and HPV-HNC cells. We first cloned several different lengths (0.17 to 1.0 kb) of the *MARCHF8* promoter regions between -840 and +160 into a luciferase reporter plasmid, pGL4.2 (**Fig. 2A**), and tested their transcription activity in HPV+ HNC (SCC152) and HPV-HNC (SCC-1) cells. Our results showed that the promoter activities of all fragments are higher in HPV+ HNC cells compared to HPV-HNC cells. Furthermore, the promoter region from -60 to -10 contains target sequences for cis-acting elements essential for *MARCHF8* promoter activity (**Fig. 2B**). Our in-silico analysis using the Eukaryotic Promoter Database (epd.epfl.ch) predicted two enhancer boxes (E-box), the known binding sites of the MYC/MAX complex, between -60 and -10 in the *MARCHF8* promoter (**Fig. 2B and Fig. S1**). To further determine if HPV oncoprotein expression increases *MARCHF8* promoter activity in HPV-HNC cells, we cotransfected SCC1 cells with the *MARCHF8* promoter-reporter construct and HPV16 E6 and/or E7 expression plasmids. Interestingly, the expression of HPV16 E6 or E6E7, but not E7 alone, significantly increases *MARCHF8* promoter activity (**Fig. 2D**). Given that both E6 and E7 expression upregulates MARCHF8 mRNA expression in N/Tert-1 cells (**Fig. 1C**), these results suggest that while E6 directly activates the *MARCHF8* promoter, E7 increases the MARCHF8 mRNA level through an alternative mechanism.

**FIG. 2.**
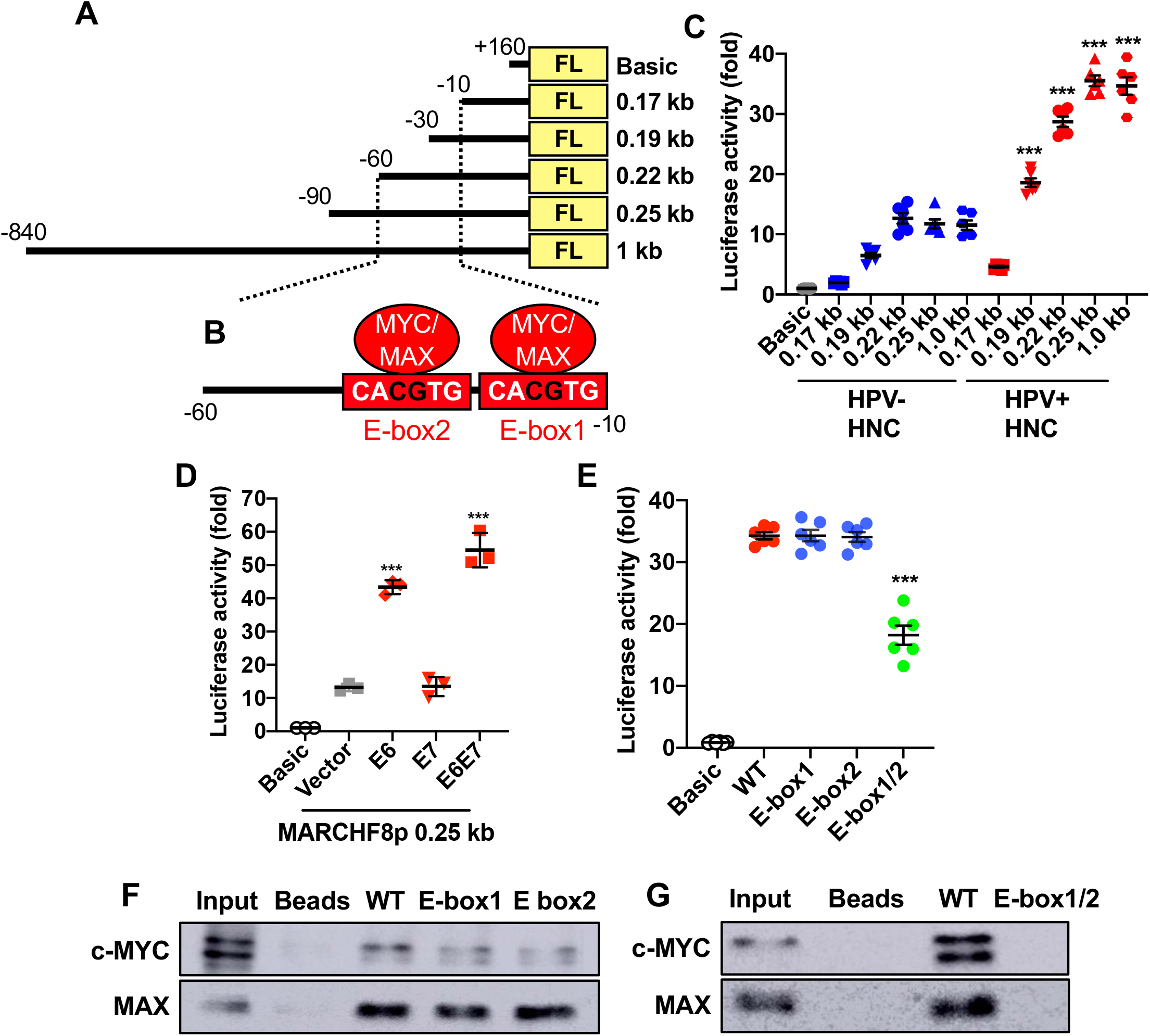
The HPV oncoprotein E6 induces the *MARCHF8* promoter activity mediated by the MYC/MAX complex. Schematic representation of the *MARCHF8* promoter regions (−840 to +160) in the firefly luciferase (FL) reporter plasmid pGL4.2 (**A**) and two E-boxes (**B**). The promoter-reporter constructs were transfected into HPV+ (SCC152) and HPV-(SCC1) cells (**C**) and cotransfected into SCC-1 HNC cells with HPV16 E6, E7, or E6 and E7 expression plasmids (**D**). SCC-152 cells were transfected with the 0.25 kb promoter (−90 to +160) reporter constructs of wildtype (WT) and E-box mutants containing single or double CG deletion in E-box1, E-box-2, and E-box1/2 (**B** and **E**). Luciferase activity was measured 48 h post transfection. Representative data of three independent experiments are shown as a fold change relative to the empty pGL4.2 vector (Basic). Shown are representative data of three repeats. *P* values were determined by Student’s *t*-test. \**p* < 0.05, \*\**p* < 0.01, \*\*\**p* < 0.001. A DNA-protein pulldown assay was performed using biotinylated 90 bp oligonucleotides containing wildtype or E-boxes mutants (−85 to +5) incubated with nuclear extracts from SCC-152 cells. The DNA-bound proteins were analyzed using anti-c-MYC and MAX antibodies (**F** and **G**) by western blotting.

To identify the cellular factors that bind to the essential *MARCHF8* promoter regions (−60 to -10) in the context of HPV16 E6, we performed a DNA-protein pulldown assay using biotinylated 90 bp oligonucleotides containing the promoter sequence between -85 and +5. Our liquid chromatography-mass spectrometry identified 83 *MARCHF8* promoter binding proteins (**Table S1**). As predicted in our in-silico analysis (**Fig. 2C and S1**), MYC-associated factor X (MAX), a member of the MYC family of transcription factors, was identified as a *MARCHF8* promoter binding protein. The MYC/MAX heterodimer complex binds to the DNA sequence designated E-box (CACGTG). To determine if the E-boxes are necessary for *MARCHF8* promoter activity, we generated a pGL4.2 reporter plasmid with a mutant 0.25 kb promoter region by deleting CG in either or both E-boxes (**Figure 2B**) and determined the promoter activity. While CG deletion in only one of the two E boxes did not change any promoter activity, CG deletions in both E-boxes significantly abrogated the *MARCHF8* promoter (**Fig. 2E**). To further validate the binding of MYC and MAX proteins to the E-boxes in the *MARCHF8* promoter, *MARCHF8* promoter binding proteins were pulled down using biotinylated 90 bp oligonucleotides (−85 to +5) containing the wild type or mutant E-boxes. Then, MYC and MAX proteins were detected by western blotting. Consistent with its promoter activity (**Fig. 2E**), while the 90 bp oligonucleotide fragment with CG deletion in only one of the two E-boxes still binds to both MYC and MAX proteins (**Fig. 2F**), CG deletion in both E-boxes completely abrogates MYC and MAX binding to the *MARCHF8* promoter (**Fig. 2G**). HPV E6 is known to interact with and stabilize the MYC/MAX complex to activate the hTERT promoter (26, 27). Thus, our findings suggest that HPV E6-induced MYC/MAX plays an important role in host transcriptional regulations, including *MARCHF8*.

### Expression of death receptors on HPV+ HNC cells is post-transcriptionally downregulated by HPV oncoproteins

MARCHF8 is known to target several membrane proteins for degradation through ubiquitination. A previous study reported that TRAIL-R1, a TNFRSF death receptor, is ubiquitinated by MARCHF8 (21). Thus, we first determined total protein levels of the TNFRSF death receptors, FAS, TRAIL-R1, and TRAIL-R2, in HPV+ HNC (SCC2, SCC90, and SCC152) and HPV-HNC (SCC1, SCC9, and SCC19) cells comparing to N/Tert-1 cells by western blotting. The results showed that FAS, TRAIL-R1, and TRAIL-R2 protein levels are significantly lower in all HPV+ HNC cells, except TRAIL-R1 in SCC2, compared to N/Tert-1 cells. In contrast, HPV-HNC cells did not show any significant changes in FAS and TRAIL-R1 except in SCC9 cells, while TRAIL-R2 showed consistent downregulation in both HPV+ and HPV-HNC cells (**Fig. 3A and 3B**). Next, cell surface expression of FAS, TRAIL-R1, and TRAIL-R2 on the HPV+ HNC and HPV-HNC cells was determined by flow cytometry. Consistent with the total protein levels, FAS, TRAIL-R1, and TRAIL-R2 expression on all HPV+ HNC cells, except TRAIL-R1 on SCC2 cells, is significantly decreased compared to N/Tert-1 cells (**Fig. 3C - 3H**). In contrast, across the HPV-HNC cells, all three death receptors are expressed at variable levels showing no clear trend of surface expression compared to N/Tert-1 cells (**Fig. 3C - 3H**). Next, we measured mRNA levels of FAS, TRAIL-R1, and TRAIL-R2 in HPV+ and HPV-HNC cells along with N/Tert-1 cells by RT-qPCR. The results showed that FAS mRNA levels are upregulated in HPV+ HNC cells, while FAS mRNA levels are not changed or slightly decreased in HPV-HNC cells compared to N/Tert-1 cells (**Fig. S1A**). In addition, mRNA expression of TRAIL-R1 (**Fig. S1B**) and TRAIL-R2 (**Fig. S1C**) is decreased in HPV-HNC cells and variable in HPV+ HNC cells. These results suggest that the downregulation of FAS, TRAIL-R1, and TRAIL-R2 expression on HPV+ HNC cells is likely caused by post-transcriptional regulation.

**FIG. 3.**
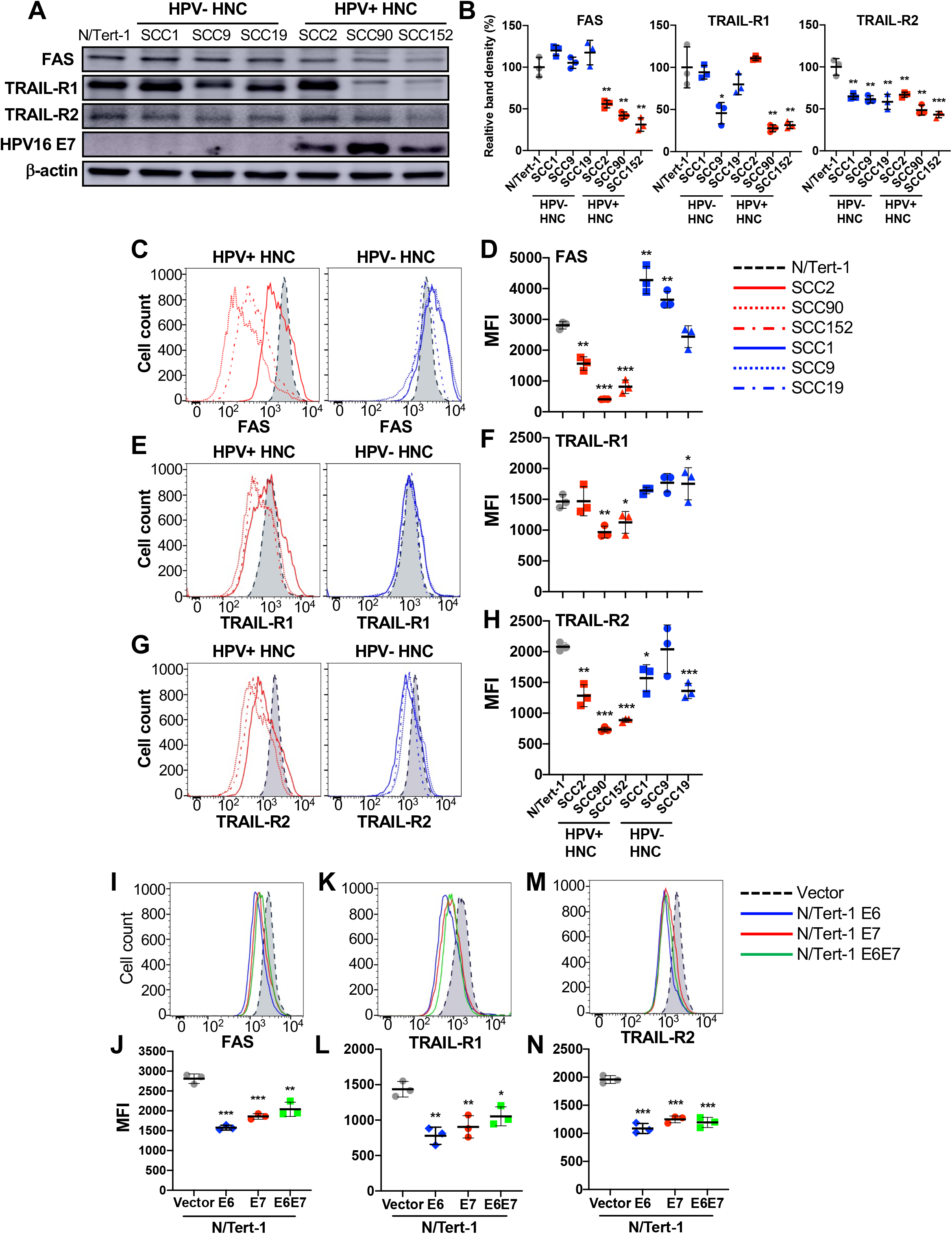
Expression of FAS, TRAIL-R1, and TRAIL-R2 is downregulated in HPV+ HNC cells. Total protein expression of FAS, TRAIL-R1, and TRAIL-R2 in HPV-(SCC1, SCC9, and SCC19) and HPV+ (SCC2, SCC90, and SCC152) HNC cells were determined by western blotting (**A**). Relative band density was quantified using NIH ImageJ (**B**). HPV16 E7 and β-actin were used as viral and internal controls, respectively. Cell surface expression of FAS (**C** and **D**), TRAIL-R1 (**E** and **F**), and TRAIL-R2 (**G** and **H**) proteins on HPV+ (SCC2, SCC90, and SCC152) and HPV-(SCC1, SCC9, and SCC19) HNC cells was analyzed by flow cytometry. Cell surface expression of FAS (**I** and **J**), TRAIL-R1 (**K** and **L**), and TRAIL-R2 (**M** and **N**) proteins on N/Tert-1 cells expressing HPV16 E6, E7, and E6E7 was analyzed by flow cytometry. Mean fluorescence intensities (MFI) of three independent experiments are shown (**D, F, H, J, L**, and **N**). All experiments were repeated at least three times, and the data shown are means ± SD. *P* values were determined by Student’s *t*-test. \**p* < 0.05, \*\**p* < 0.01, \*\*\**p* < 0.001.

Next, we determined whether the HPV16 oncoproteins E6 and E7 contribute to the downregulation of FAS, TRAIL-R1, and TRAIL-R2, using N/Tert-1 cells expressing E6 (N/Tert-1 E6), E7 (N/Tert-1 E7), or both E6 and E7 (N/Tert-1 E6E7). Interestingly, our data showed that expression of either or both HPV16 E6 and E7 is sufficient for a significant decrease in FAS, TRAIL-R1, and TRAIL-R2 expression on N/Tert-1 cells (**Fig. 3I - 3N**). Our results suggest that surface expression of the TNFRSF death receptors on HPV+ HNC cells is post-transcriptionally downregulated by the HPV oncoproteins E6 and E7.

### Knockdown of *MARCHF8* expression increases FAS, TRAIL-R1, and TRAIL-R2 expression in HPV+ HNC cells

To determine if the *MARCHF8* upregulation by E6 and E7 is responsible for the downregulation of FAS, TRAIL-R1, and TRAIL-R2 in HPV+ HNC cells, we knocked down *MARCHF8* expression in SCC152 cells using five unique shRNAs against *MARCHF8* (shR-MARCHF8, clones 1-5) delivered by lentiviruses and selected by puromycin treatment. All five shR-MARCHF8s in SCC152 cells showed a ∼50% decrease in total MARCHF8 protein levels compared to the control SCC152 cells with nonspecific scrambled shRNA (shR-scr) (**Fig. 4A and 4B**). Using western blotting and flow cytometry, we analyzed protein expression of FAS, TRAIL-R1, and TRAIL-R2. We found that both total (**Fig. 4A and 4B**) and cell surface (**Fig. 4C-4H**) protein levels of FAS, TRAIL-R1, and TRAIL-R2 are significantly increased by all five cell lines with *MARCHF8* knockdown. We also knocked down *MARCHF8* expression in another HPV+ HNC cell line, SCC2, using three shRNAs and confirmed the increase of total (**Fig S2A and S2B**) and surface (**Fig S2C - S2H**) FAS, TRAIL-R1, and TRAIL-R2 expression by *MARCHF8* knockdown. To determine whether knockdown of *MARCHF8* expression affects mRNA expression of FAS, TRAIL-R1, and TRAIL-R2, we measured mRNA levels of FAS, TRAIL-R1, and TRAIL-R2 in the SCC152 and SCC2 cells with *MARCHF8* knockdown by RT-qPCR. The results showed no significant changes in FAS mRNA expression by *MARCHF8* knockdown in both SCC152 (**Fig. S3A**) and SCC2 cells (**Fig. S3B**). Additionally, mRNA expression of TRAIL-R1 and TRAIL-R2 is decreased in SCC152 cells (**Fig. S3B and S3C**) but not in SCC-2 cells (**Fig. S3E and S3F**) with *MARCHF8* knockdown, indicating that the decrease of FAS, TRAIL-R1, and TRAIL-R2 protein levels in HPV+ HNC cells are not caused by a reduction in their mRNA levels. These results suggest that HPV oncoprotein-induced MARCHF8 post-transcriptionally downregulates the TNFRSF death receptors.

**FIG. 4.**
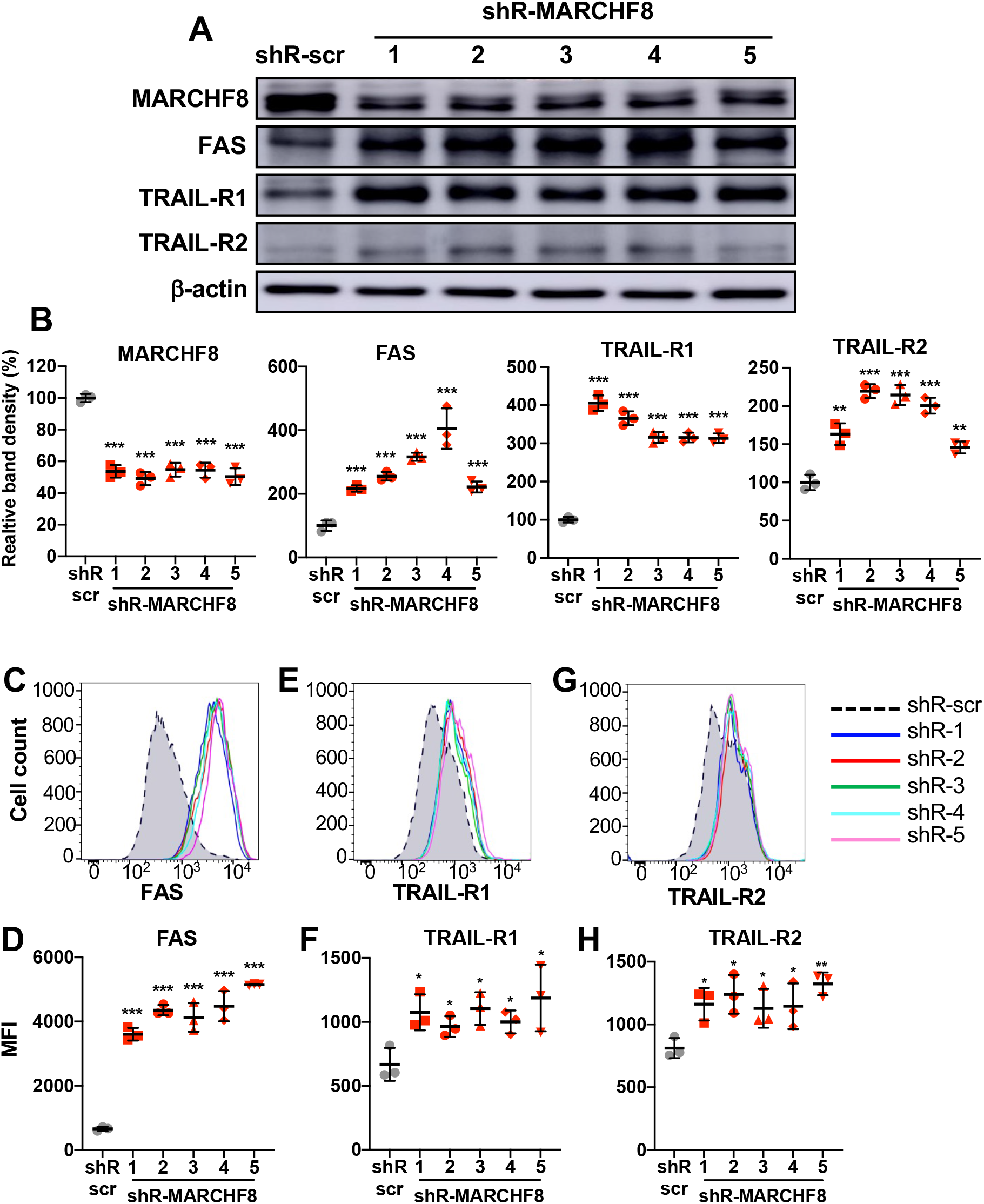
Knockdown of *MARCHF8* expression increases FAS, TRAIL-R1, and TRAIL-R2 protein expression in HPV+ HNC cells. HPV+ HNC (SCC152) cells were transduced with one of five lentiviral shRNAs against *MARCHF8* (shR-MARCHF8 clones 1-5) or scrambled shRNA (shR-scr) as a control. Protein expression of MARCHF8, FAS, TRAIL-R1, and TRAIL-R2 was determined by western blotting (**A**). Relative band density was quantified using NIH ImageJ (**B**). β-actin was used as an internal control. The data shown are means ± SD of three independent experiments. Cell surface expression of FAS (**C** and **D**), TRAIL-R1 (**E** and **F**), and TRAIL-R2 (**G** and **H**) proteins were analyzed by flow cytometry. Mean fluorescence intensities (MFI) of three independent experiments are shown (**D, F**, and **H**). *P* values were determined by Student’s *t*-test. \**p* < 0.05, \*\**p* < 0.01, \*\*\**p* < 0.001.

### MARCHF8 protein interacts with and ubiquitinates FAS, TRAIL-R1, and TRAIL-R2 proteins in HPV+ HNC cells

Previous studies have shown that MARCHF8, a membrane-associated ubiquitin ligase, binds to and ubiquitinates several membrane receptor proteins for degradation (21, 28, 29). Thus, we hypothesized that MARCHF8 upregulated by the HPV oncoproteins targets FAS, TRAIL-R1, and TRAIL-R2 proteins for ubiquitination and degradation. First, to determine if MARCHF8 protein binds to FAS, TRAIL-R1, and TRAIL-R2, we pulled down MARCHF8 protein in whole cell lysates from SCC152 cells treated with the proteasome inhibitor MG132 using magnetic beads conjugated with an anti-MARCHF8 antibody. The western blot analyses showed that FAS, TRAIL-R1, and TRAIL-R2 proteins were detected in the MARCHF8 protein complex pulled down with an anti-MARCHF8 antibody (**Fig. 5A**). Reciprocally, the co-immunoprecipitation of FAS protein in the same whole cell lysates using an anti-FAS antibody showed MARCHF8 protein (**Fig. 5B**). Next, to determine if MARCHF8 induces ubiquitination of FAS, TRAIL-R1, and TRAIL-R2 proteins, we pulled down ubiquitinated proteins in whole-cell lysates from SCC152 cells treated with MG132 using magnetic beads conjugated with an anti-ubiquitin antibody. The results showed that the levels of ubiquitinated FAS (**Fig. 5C**), TRAIL-R1 (**Fig. 5D**), and TRAIL-R2 (**Fig. 5E**) proteins were decreased in SCC152 cells by *MARCHF8* knockdown, despite the significantly higher levels of total input proteins of FAS, TRAIL-R1 and TRAIL-R2, compared to SCC152 cells with shR-scr. These results suggest that HPV-induced MARCHF8 binds to and ubiquitinates the TNFRSF death receptors in HPV+ HNC cells.

**FIG. 5.**
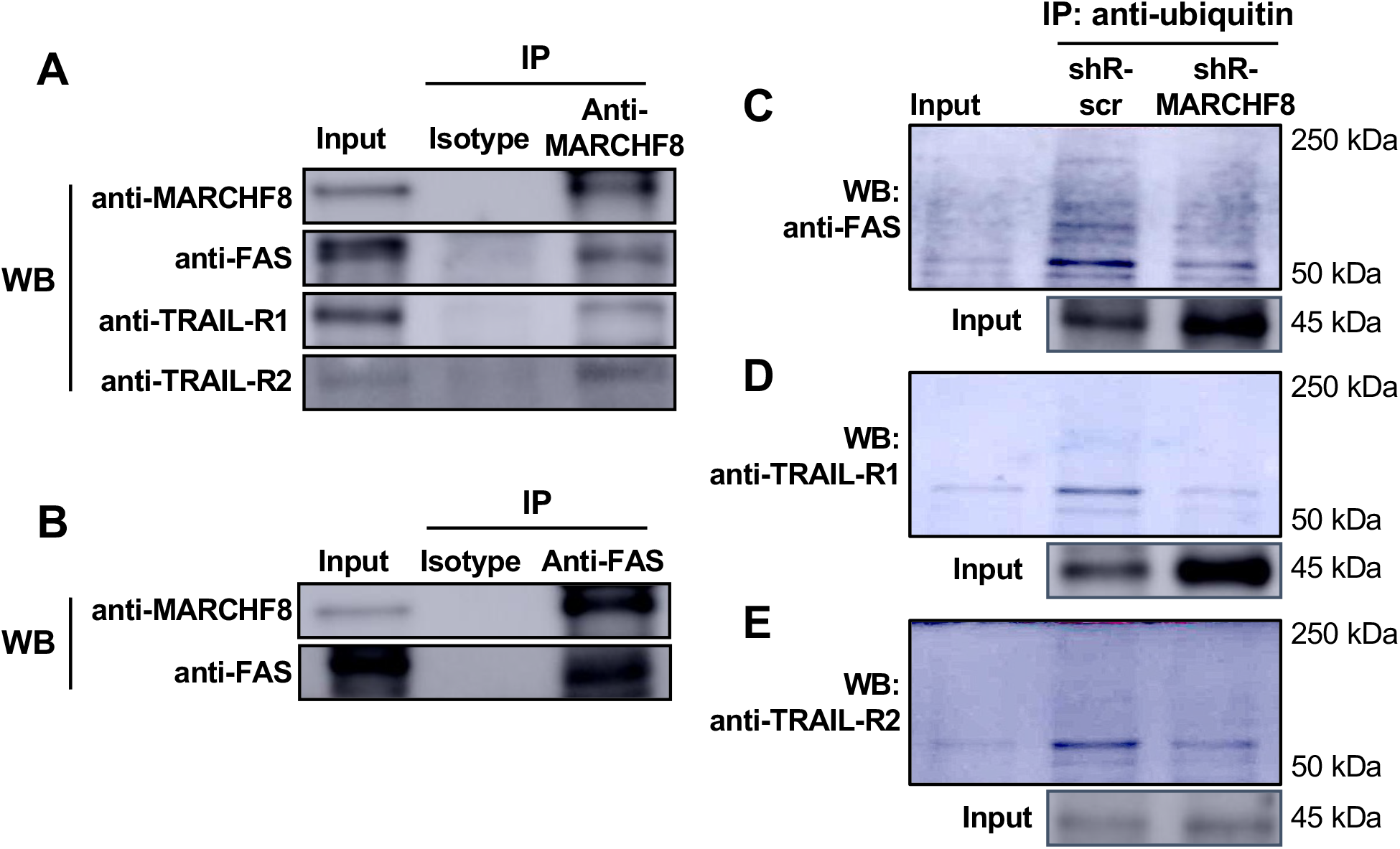
MARCHF8 protein interacts with and ubiquitinates FAS, TRAIL-R1, and TRAIL-R2 proteins. MARCHF8 (**A**) and FAS (**B**) were pulled down from the cell lysate of HPV+ HNC (SCC-152) cells treated with a proteasome inhibitor MG132 using anti-MARCHF8 (**A**) and anti-FAS (**B**) antibodies, respectively. Western blotting detected FAS, TRAIL-R1, TRAIL-R2, and MARCHF8 proteins in the immunoprecipitated proteins. Ubiquitinated proteins were pulled down from the cell lysate of HPV+ HNC (SCC152) cells with scrambled shRNA (shR-scr) or shRNA against MARCHF8 (shR-MARCHF8 clone 3) treated with a proteasome inhibitor MG132 using an anti-ubiquitin antibody (**C** - **E**). FAS (**C**), TRAIL-R1 (**D**), and TRAIL-R2 (**E**) proteins were detected in the immunoprecipitated proteins by western blotting. All experiments were repeated at least three times.

### Knockdown of *MARCHF8* expression increases apoptosis of HPV+ HNC cells

Since expression of the death receptors is significantly upregulated by *MARCHF8* knockdown (**Fig. 4 and S2**), we hypothesized that the HPV oncoproteins inhibit host cell apoptosis by inducing MARCHF8-mediated degradation of FAS, TRAIL-R1, and TRAIL-R2 proteins. To test this hypothesis, we determined whether knockdown of *MARCHF8* expression in HPV+ HNC cells enhances FAS-mediated apoptosis. SCC152 and SCC2 cells with two shR-MARCHF8s or shR-scr were first sensitized for apoptosis by treating with the soluble recombinant FAS ligand (rFAS-L), and apoptotic cells were quantified by detecting annexin V- and 7-aminoactinomycin D (7-AAD)-positive cells. The results showed that both SCC152 (**Fig 6A and 6B**) and SCC2 (**Fig 6C and 6D**) cells with *MARCHF8* knockdown displayed significantly increased percentages of apoptotic cells compared to the corresponding cells with shR-scr under the sensitization with rFAS-L. We also performed the apoptosis assay by sensitizing SCC152 and SCC2 cells with an anti-FAS antibody and confirmed that knockdown of *MARCHF8* significantly increases apoptosis of HPV+ HNC cells (data not shown). These results suggest that by ubiquitinating and degrading FAS proteins, MARCHF8, transcriptionally upregulated by HPV oncoproteins, inhibits host cell apoptosis.

**FIG. 6.**
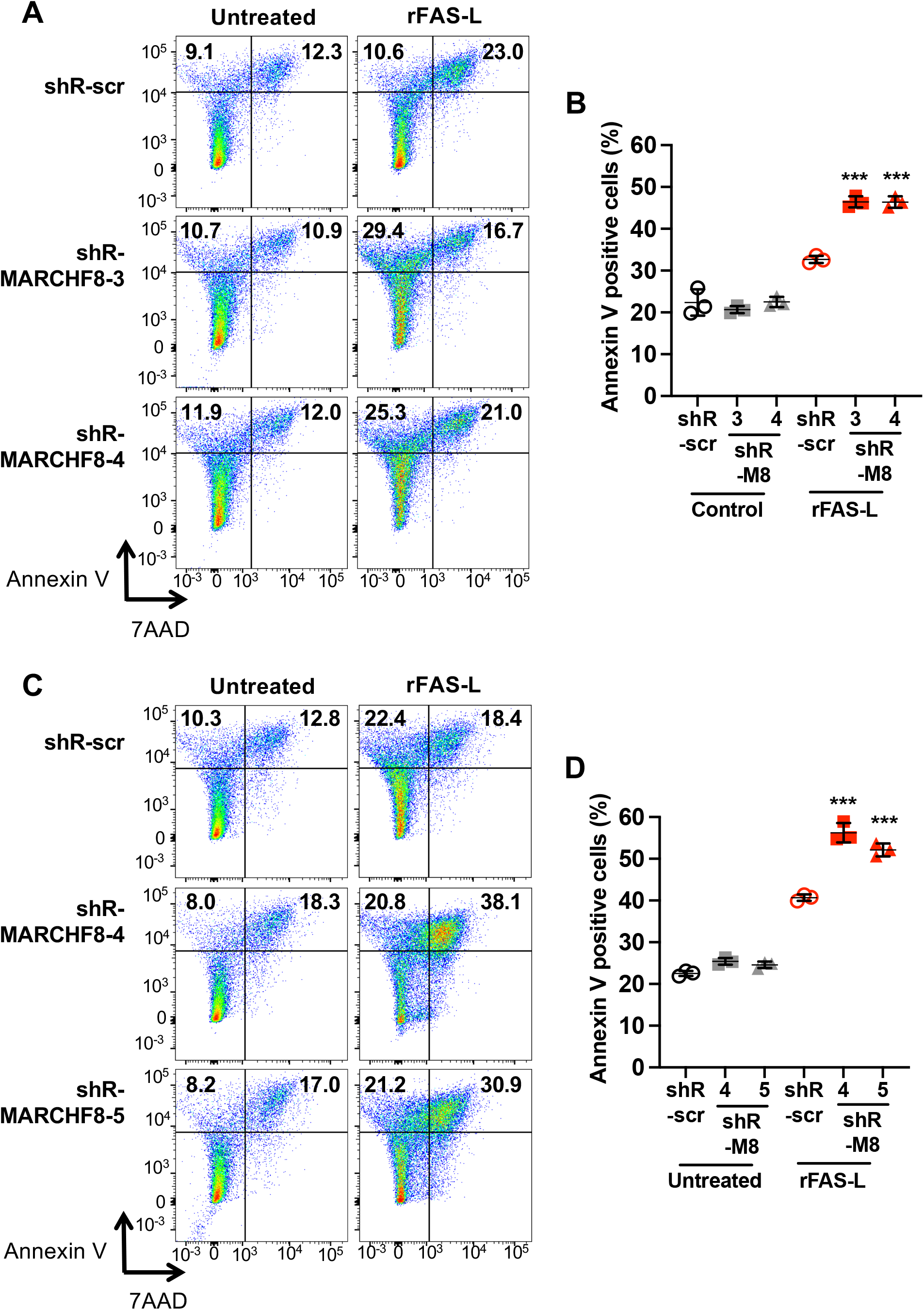
Knockdown of *MARCHF8* enhances apoptosis of HPV+ HNC cells. Two HPV+ HNC cells, SCC152 (**A** and **B**) and SCC2 (**C** and **D**), with scrambled shRNA (shR-scr) or two shRNAs against MARCHF8 (shR-MARCHF8), were treated with the recombinant FAS ligand (rFAS-L). The cells were stained with an anti-annexin V antibody and 7-AAD and analyzed by flow cytometry. The percentage of cells with positive staining is indicated (**A** and **C**). The data shown are means ± SD of three independent experiments (**B** and **D**). *P* values were determined by Student’s *t*-test. \*\*\**p* < 0.001.

### *Marchf8* expression is upregulated, and death receptor expression is downregulated in HPV+ mouse oral cancer cells

To investigate the inhibition of apoptosis by HPV-induced MARCHF8 degradation of the TNFRSF death receptors, we adopted a mouse model of HPV+ HNC with *Hras*-transformed mouse oral epithelial cells expressing HPV16 E6 and E7 (MOE/E6E7 or mEERL) that form tumors in immunocompetent syngeneic C57BL/6 mice (30). First, we determined total protein levels of MARCHF8, FAS, TRAIL-R1, and TRAIL-R2 in mEERL cells compared to normal immortalized mouse oral epithelial (NiMOE) cells and mouse MOE cells transformed with *Hras* and shR-Ptpn14 (HPV-MOE). The results from the mouse oral cancer cells were consistent with those from human HNC cells showing a significant increase in MARCHF8 protein levels and a decrease in FAS, TRAIL-R1, and TRAIL-R2 protein levels in mEERL cells compared to NiMOE cells (**Fig. 7A and 7B**). Similarly, surface expression of FAS and TRAIL-R2 proteins was significantly downregulated in mEERL cells compared to NiMOE cells (**Fig. 7C - 7F**). While there were no significant changes in TRAIL-R2 protein levels in HPV-MOE cells compared to NiMOE cells (**Fig. 7A, 7B, 7E, and 7F**), total and cell surface expression of FAS and TRAIL-R1 proteins were decreased in HPV-MOE cells compared to NiMOE cells (**Fig. 7B - 7D**). These results suggest that mEERL cells recapitulate our findings from human HPV+ HNC cells that HPV-induced MARCHF8 degrades the TNFRSF death receptors.

**FIG. 7.**
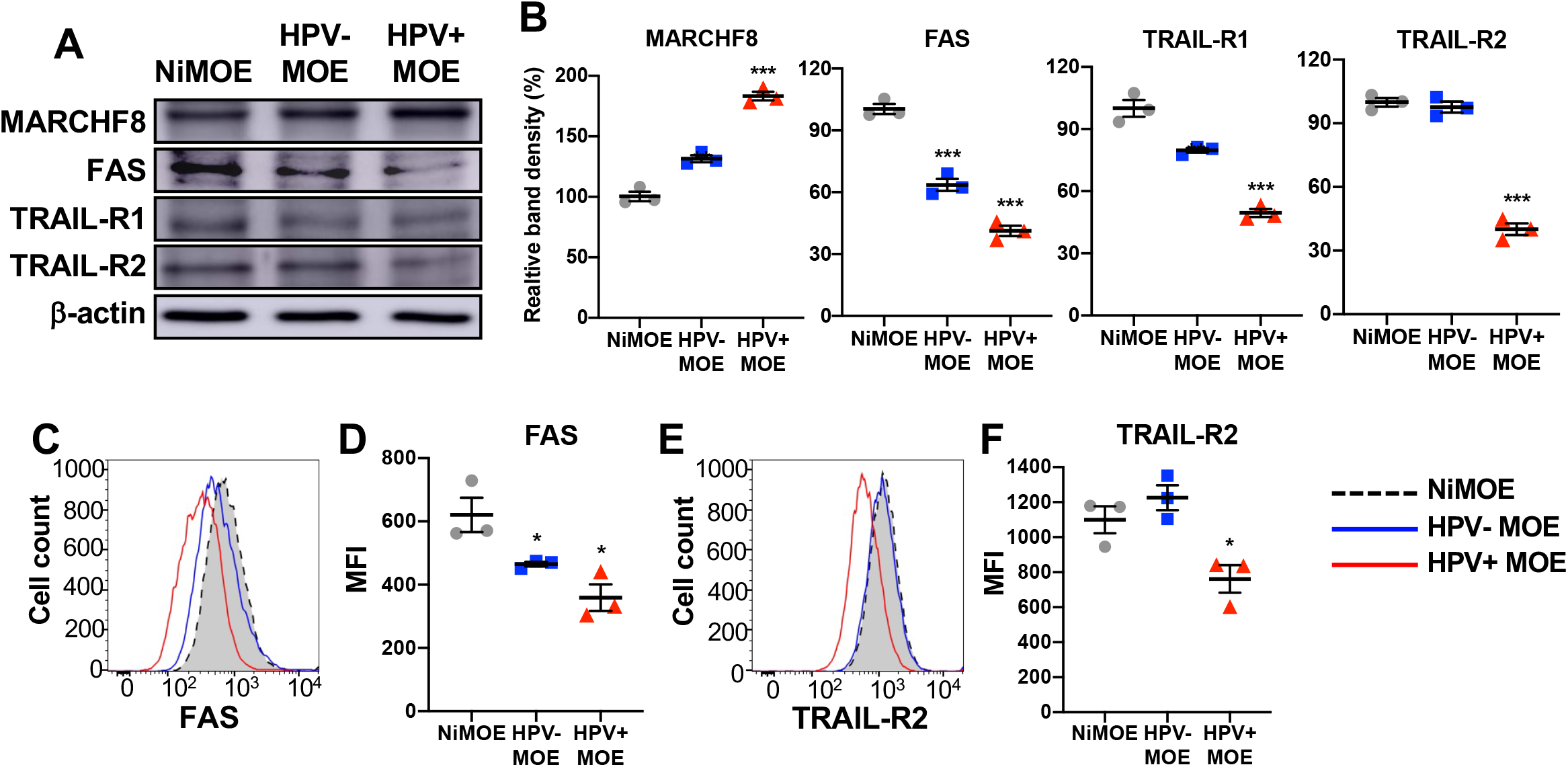
*MARCHF8* is upregulated, and death receptor expression is downregulated in HPV+ mouse oral cancer cells. Mouse MARCHF8, FAS, TRAIL-R1, and TRAIL-R2 protein levels in mouse normal immortalized (NiMOE), HPV-transformed (HPV-MOE), and HPV+ transformed (HPV+ MOE) oral epithelial cells were determined by western blotting (**A**). β-actin was used as a loading control. The relative band density was quantified using NIH ImageJ (**B**). Cell surface expression of FAS (**C and D**) and TRAIL-R2 (**E and F**) proteins on NiMOE (dotted black line), HPV-MOE (blue line), and HPV+ MOE (red line) cells were analyzed by flow cytometry. Mean fluorescence intensities (MFI) of three independent experiments are shown (**D** and **F**). All experiments were repeated at least three times, and the data shown are means ± SD. *P* values were determined by Student’s *t*-test. \**p* < 0.05, \*\*\**p* < 0.001.

### *Marchf8* knockout in HPV+ mouse oral cancer cells restores FAS, TRAIL-R1, and TRAIL-R2 expression and enhances apoptosis

To determine if the high levels of *Marchf8* expression in HPV+ MOE cells are responsible for the downregulation of FAS, TRAIL-R1, and TRAIL-R2, we established *Marchf8* knockout mEERL (mEERL/*Marchf8*^-/-^) cell lines using lentiviral transduction of Cas9 and three small guide RNAs (sgRNAs) against *Marchf8* (sgR-*Marchf8*, clones 1-3). mEERL cells transduced with Cas9 and scrambled sgRNA (mEERL/scr) were used as a control. mEERL cell lines transduced with two (clones 2 and 3) of the three sgR-*Marchf8*s showed a ∼75% decrease in *Marchf8* expression compared to mEERL/scr cells (**Fig. 8A and 8B**). Consistent with the data from human HPV+ HNC cells presented in **Fig. 4**, mEERL/*Marchf8*^-/-^ cells showed significantly upregulated total (**Fig. 8A and 8B**) and surface expression (**Fig. 8C - 8F**) of FAS and TRAIL-R2 compared to mEERL/scr cells. Next, we determined whether *Marchf8* knockout enhances apoptosis of mEERL/*Marchf8*^-/-^ cells by sensitizing with rFAS-L as described in **Fig. 6**. The results showed that mEERL/*Marchf8*^-/-^ cells showed significantly increased annexin V- and 7-AAD-positive cells compared to mEERL/scr cells (**Fig. 8G and 8H**). These results are consistent with our findings in human HPV+ HNC that HPV-induced *MARCHF8* expression inhibits host cell apoptosis by degrading the TNFRSF death receptors.

**FIG. 8.**
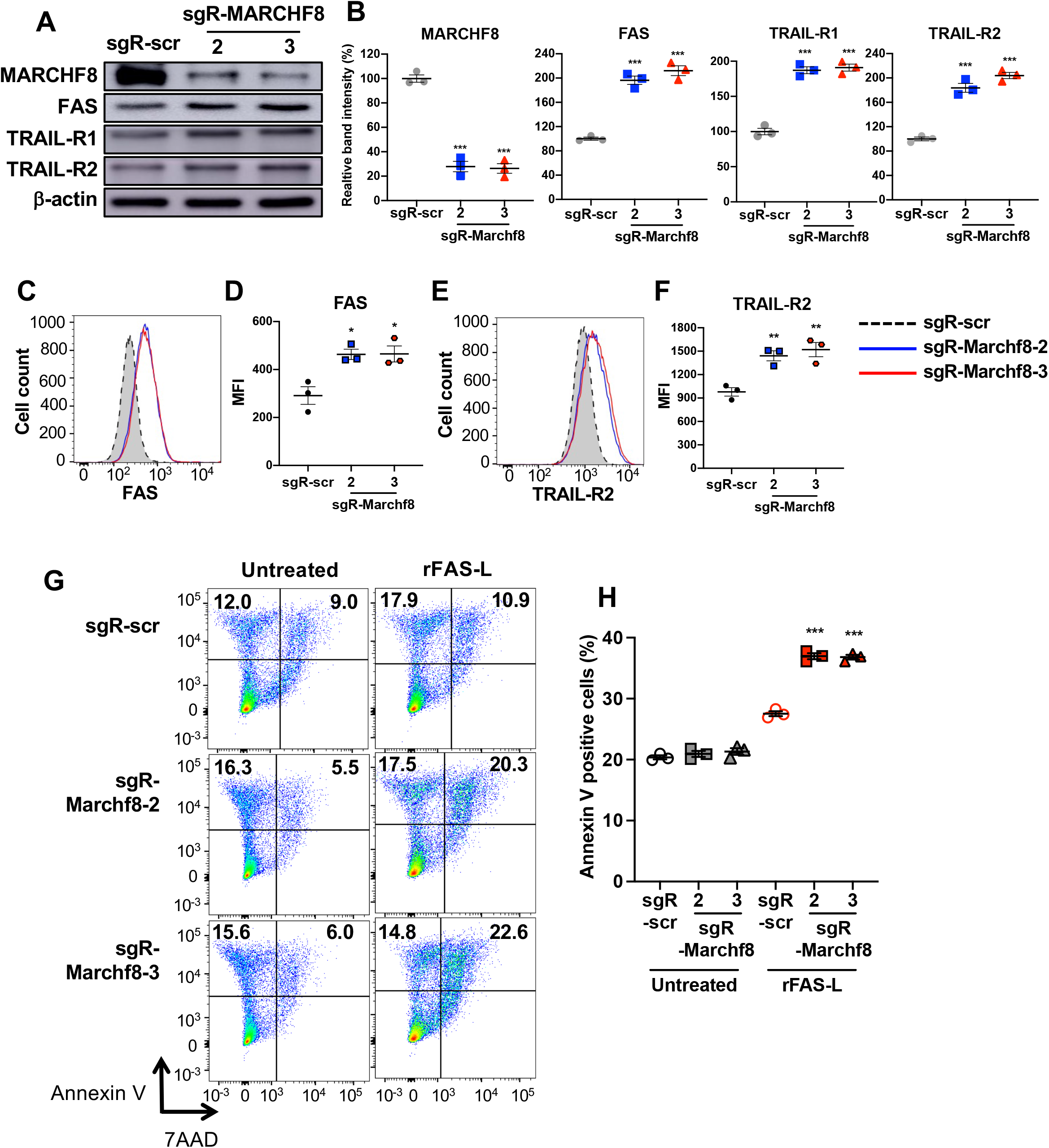
Knockout of *Marchf8* expression increases FAS, TRAIL-R1, and TRAIL-R2 protein levels and enhances apoptosis of HPV+ mouse oral cancer cells. mEERL cells were transduced with lentiviral Cas9 and one of two sgRNAs against Marchf8 (sgR-Marchf8-2 and sgR-Marchf8-3) or scrambled sgRNA (sgR-scr). Protein levels of MARCHF8, FAS, TRAIL-R1, and TRAIL-R2 were determined by western blotting (**A**). Relative band density was quantified using NIH ImageJ (**B**). β-actin was used as a loading control. The data shown are means ± SD of three independent experiments. Cell surface expression of FAS (**C** and **D**) and TRAIL-R2 (**E** and **F**) proteins were analyzed by flow cytometry. Mean fluorescence intensities (MFI) of three independent experiments are shown (**D** and **F**). *P* values were determined by Student’s *t*-test. \**p* < 0.05, \*\**p* < 0.01, \*\*\**p* < 0.001. Untreated and rFAS-L-treated mEERL cells (**G**) with sgR-Marchf8-2, sgR-Marchf8-3, or sgR-scr were stained with an anti-annexin V antibody and 7-AAD and analyzed by flow cytometry. The percentage of cells with positive staining is indicated (**G**). The data shown are means ± SD of three independent experiments (**H**). *P* values were determined by Student’s *t*-test. \*\*\**p* < 0.001.

### *Marchf8* knockout in HPV+ HNC cells suppresses tumor growth in vivo

To determine whether *Marchf8* knockout in HPV+ HNC suppresses tumor growth in vivo, we injected syngeneic C57BL/6J mice subcutaneously with either of two 5 × 10^5^ of mEERL/*Marchf8*^-/-^ cell lines or mEERL/scr cells into the flank. Tumor growth was monitored twice a week for 12 weeks. All ten mice injected with mEERL/scr cells showed vigorous tumor growth (**Fig. 9A and 9C**) and succumbed to tumor burden in ∼7 weeks post injection (**Fig. 9B**). In contrast, only two out of ten mice each injected with either of two mEERL/*Marchf8*^-/-^ cell lines showed robust tumor growth (**Fig. 9A, 9D, and 9E**) and died in 8 weeks post injection (**Fig. 9B**). Further, no tumor formation was observed in three and one mice injected with mEERL/*Marchf8*^-/-^ cell lines, sgR-Marchf8-2 and sgR-Marchf8-3, respectively, over 12 weeks post injection (**Fig. 9D and 9E**). Our results suggest that MARCHF8 is a potent tumor promoter that plays an important role in cancer progression by inducing the degradation of the TNFRSF death receptors and blocking cell apoptosis.

**FIG. 9.**
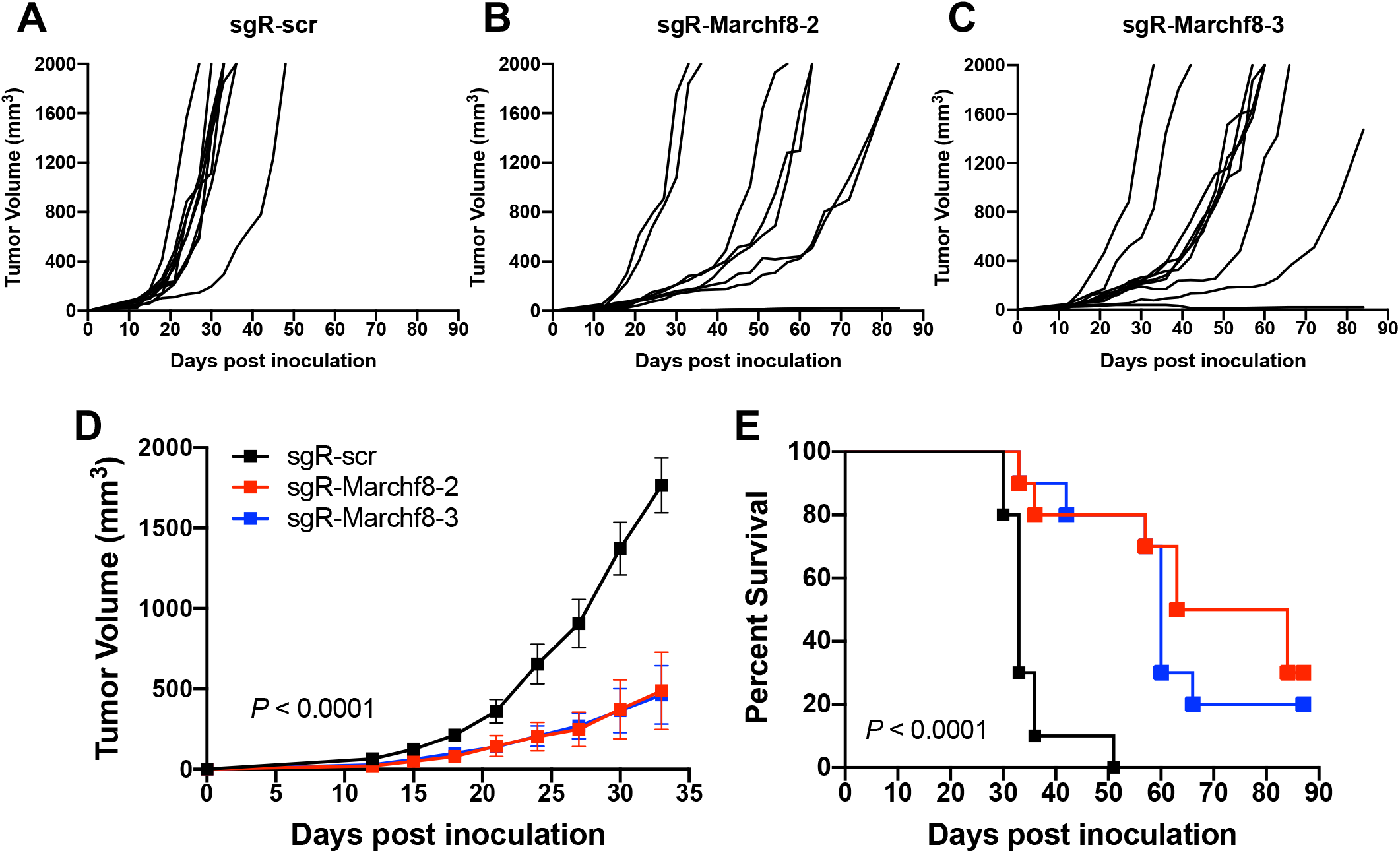
Knockout of *Marchf8* expression suppresses HPV+ HNC tumor growth in vivo. *Marchf8*-knockout mEERL cells (mEERL/*Marchf8*^-/-^) were generated by lentiviral Cas9, and two sgRNAs targeting *Marchf8* (sgR-Marchf8-2 and sgR-Marchf8-3) or scrambled sgRNA (sgR-scr). mEERL/scr (**A**) or mEERL/*Marchf8*^-/-^ (**B** and **C**) cells were injected into the rear right flank of C57BL/6J mice (*n* = 10 per group). Tumor volume was measured twice a week (**A**-**D**). Survival rates of mice were analyzed using a Kaplan-Meier estimator (**E**). The time to event was determined for each group, with the event defined as a tumor size larger than 2000 mm^3^. The data shown are means ± SD. *P* values of mice injected with mEERL/*Marchf8*^-/-^ cells compared with mice injected with mEERL/scr cells were determined for tumor growth (**D**) and survival (**E**) by two-way ANOVA analysis. Shown are representative of two independent experiments.

**FIG. 10.**
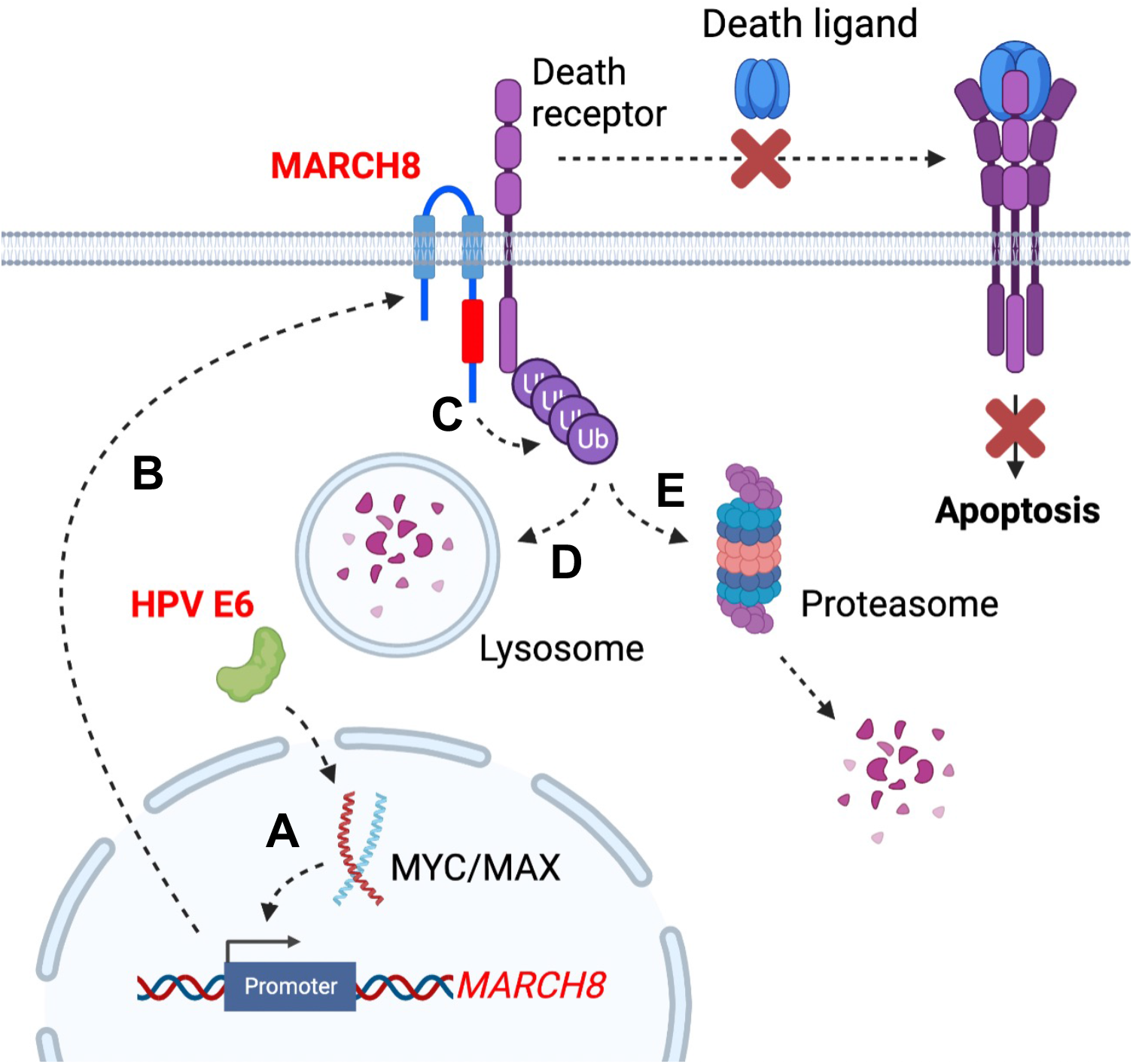
The schematic diagram summarizes that HPV E6-induced *MARCHF8* expression inhibits host cell apoptosis by degrading the TNFRSF death receptors. The HPV oncoprotein E6 activates the *MARCHF8* promoter activity through the MYC/MAX transcription factor complex (**A**) and upregulates cell surface expression of the MARCHF8 protein (**B**). MARCHF8 protein binds to and ubiquitinates the TNFRSF death receptors FAS, TRAIL-R1, and TRAIL-R2 (**C**). Ubiquitinated TNFRSF death receptors may be degraded by lysosomes (**D**) or proteasomes in the cytoplasm (**E**).

## DISCUSSION

As cancer cells must prevent apoptosis for survival, the TNFRSF death receptors are frequently dysregulated in cancer cells (31). To inhibit death receptor-mediated apoptosis, cancer cells often repress *FAS* expression by histone modification (32), polymorphism (33), and hypermethylation of the promoter region (34, 35). Cancer cells also abrogate the function of the death receptors by generating loss-of-function mutations (36, 37) and inducing the expression of inhibitory molecules such as FLICE-like inhibitory protein (38–40), decoy receptors and ligands (41, 42), and microRNA miR-196b (43, 44). Additionally, cancer cells dysregulate the trafficking of death receptors to interfere with their cell surface expression (45, 46). Interestingly, MARCHF8 ubiquitinates TRAIL-R1 and diminishes its cell surface expression on breast cancer cells (21). In addition, a previous study has shown that proteasome inhibitor treatment upregulates TRAIL-R2 protein and induces apoptosis in prostate cancer cells (47). These findings suggest that inhibiting TNFRSF death receptors-mediated apoptosis through ubiquitination plays an important role in cancer cell survival. Nevertheless, the detailed mechanisms by which HPV+ HNC cells inhibit TNFRSF death receptors were mostly unknown.

This study reports that MARCHF8, an E3 ubiquitin ligase transcriptionally activated by the HPV oncoprotein E6 and the MYC/MAX complex, downregulates surface expression of FAS, TRAIL-R1, and TRAIL-R2 on HPV+ HNC cells. The TNFSFR death receptors, characterized by a cytoplasmic tail termed the death domain, play a crucial role in apoptosis upon their interactions with specific extracellular ligands such as FAS-L and TRAIL (a.k.a. TNFSF10) (48–50). FAS, the first member of the TNFRSF expressed in various tissues, interacts with FAS-L (a.k.a. CD178) to trigger apoptosis (51, 52). TRAIL-R1 and TRAIL-R2 interact with TRAIL (a.k.a. TNFSF10) (53, 54). In contrast, the other TRAIL-Rs, TRAIL-R3, and TRAIL-R4 do not have any death domain and are not considered death receptors.

MARCHF8, originally named cellular MIR (c-MIR), was first discovered as a human homolog of two KSHV proteins, a modulator of immune recognition 1 and 2 (MIR1 and MIR2, a.k.a. K3 and K5, respectively) (14). Similar to KSHV MIR1 and MIR2, MARCHF8 downregulates the surface expression of various immunoreceptors, including MHC-II (55), CD44 (20, 56), CD81(56), and CD86 (57). MARCHF8 also decreases the surface expression of TRAIL-R1 (21, 58) and inhibits apoptosis. In addition to KSHV, many other viruses employ various strategies to restrain host cell apoptosis, which is considered an antiviral host innate response (59). For example, DNA viruses such as adenovirus encode viral proteins that downregulate death receptors and inhibit caspase activation (60). It has been suggested that some of these anti-apoptotic functions of the viruses contribute to the oncogenic process during virus-driven cancer progression(60–63).

High-risk HPVs have been shown to inhibit host cell apoptosis by targeting several different apoptotic mechanisms, especially the TNFRSF death receptors and their signaling (64). While HPV E5 is involved in impeding TNFRSF-mediated apoptosis (65), E6 plays a crucial role in the inhibition of host cell apoptosis by interacting with TNFRSF1A and FAS-associated death domain (FADD) and facilitating their degradation (12, 66). Further, knockdown of E6 expression or treatment of the proteasome inhibitor significantly enhances TNFRSF death receptor-mediated apoptosis of cervical cancer cells (67, 68). These results indicate that E6-mediated inhibition of apoptosis through TNFRSF death receptors is critical for cancer cell survival. However, the mechanism of E6-mediated degradation of the TNFRSF death receptors was mostly elusive. Our study has revealed that HPV E6 induces the degradation of the TNFRSF death receptors by activating the *MARCHF8* promoter through the interaction of the MYC/MAX complex (**Fig. 2**), a well-known oncogenic transcription factor complex that also activates hTERT transcription (27, 69). Liu et al. previously showed that MYC overexpression could replace HPV E6 to immortalize keratinocytes in the presence of HPV E7 despite the lack of p53 degradation (70). In addition, E6 stabilizes MYC by enhancing its O-linked GlcNAcylation (26), suggesting that E6-induced MYC plays an important role in E6-mediated oncogenesis. While hTERT induction by MYC is crucial, our findings indicate that *MARCHF8* activation by MYC may also play an essential role in cell immortalization by inhibiting apoptosis. However, as MYC is also involved in p53-induced apoptosis (71), the E6 function in p53 degradation may still be necessary for cancer progression. In addition, our result showed that keratinocytes expressing HPV16 E7 alone also had increased levels of MARCHF8 mRNA and protein (**Fig. 1C**). This suggests that multiple mechanisms may mediate the upregulation of MARCHF8 in HPV+ HNC.

It has been discovered that *MARCHF8* expression is upregulated in gastric and esophageal cancer (22), and its expression is associated with poor prognosis (58, 72). In addition, MARCHF8 ubiquitinates TRAIL-R1 and decreases apoptosis in gastric and breast cancer cells (21, 58), and silencing of MARCHF8 induces apoptosis and suppresses cell proliferation, invasion, and migration of cancer cells (22, 24). As described above, MARCHF8 plays a vital role in immune suppression by degrading MHC-II and CD86 (18, 19). Our study also showed that *MARCHF8* knockout dramatically suppresses tumor growth in vivo (**Fig. 9**). Together, these results suggest that MARCHF8, as a tumor promoter, could be a potential target for cancer therapy to induce cancer cell apoptosis and antitumor immune responses.

Despite these protumor functions in several cancers, including HPV+ HNC, some studies have shown potential antitumor activity of MARCHF8. *Overexpression of MARCHF8* inhibits NSCLC cell proliferation and metastasis via the PI3K and mTOR signaling pathways (72). In addition, *MARCHF8* overexpression also promotes apoptosis and hinders tumorigenesis and metastasis of breast cancer cells by downregulating CD44 and STAT3 (73). These results imply that MARCHF8 may differentially contribute to cancer development and that other MARCHF family members, such as MARCHF1, MARCHF4, and MARCHF9, may have similar functions in the place of MARCHF8. Therefore, while our study clearly shows the function of MARCHF8 as an oncoprotein in HPV+ HNC, further investigation is required, as MARCHF8 and other MARCHF family members exhibit extensive roles in the regulation of various membrane proteins involved in cellular homeostasis, apoptosis, and immune responses.

## MATERIALS AND METHODS

### Cell lines

HPV+ (SCC2, SCC90, and SCC152) and HPV-(SCC1, SCC9, and SCC19) HNC cells and 293FT cells were purchased from the American Type Culture Collection and Thermo Fisher, respectively. These cells were cultured and maintained as described (74–77). The N/Tert-1 cells (78) expressing HPV16 E6 (N/Tert-1-E6), E7 (N/Tert-1-E7), and E6 and E7 (N/Tert-1-E6E7) were previously generated (79, 80) and maintained in keratinocyte serum-free medium supplemented with epidermal growth factor (EGF), bovine pituitary extract, and penicillin/streptomycin (Thermo Fisher). The mouse oropharyngeal epithelial (MOE) cell lines, NiMOE, mEERL, and MOE/shPtpn13 were obtained from John Lee (30) and cultured in E-medium (DMEM and F12 media supplemented with 0.005% hydrocortisone, 0.05% transferrin, 0.05% insulin, 0.0014% triiodothyronine, 0.005% EGF, and 2% FBS) as previously described (81).

### Flow cytometry

Single-cell suspensions were prepared, counted, and analyzed using specific antibodies (**Table S2**) by an LSRII flow cytometer (BD Biosciences), as previously described (82). Data were analyzed using the FlowJo software (Tree Star). Apoptotic cells were detected using the PE Annexin V Apoptosis Detection Kit with 7-AAD according to the manufacturer’s protocol (BioLegend) and analyzed using an LSRII flow cytometer.

### Lentivirus production and transduction

The shRNAs targeting human *MARCHF8* were purchased from Sigma-Aldrich. The sgRNAs targeting mouse *Marchf8* were designed by the web-based software ChopChop (http://chopchop.cbu.uib.no) (83). The sgRNAs were synthesized and cloned into the lentiCRISPR v2-blast plasmid (Addgene) using ligating duplex oligonucleotides containing BsmBI restriction sites purchased from Integrated DNA Technologies (IDT). All shRNA and sgRNA sequences are listed in **Table S3 and S4**, respectively. Lentiviruses containing shRNA or sgRNA were produced using 293FT cells with packaging constructs pCMV-VSVG and pCMV-Delta 8.2 (Addgene). The lentiviruses were collected 48 hrs post transfection and concentrated by ultracentrifugation at 25,000 rpm for 2 hrs. Cells were incubated with lentiviruses for 48 hrs in the presence of polybrene (8 μg/ml) and selected with blasticidin (8 μg/ml).

### DNA-protein pulldown assay

The assay was performed as previously described (84). Briefly, biotinylated 90 bp oligonucleotides containing the *MARCHF8* promoter sequence between -85 and +5 (wild type or the E-box mutants) (**Fig. 2 and Fig. S1**) were synthesized and incubated with 100 μl of M-280 streptavidin-coated magnetic beads (Dynal) for 1 h. The beads were collected using a magnetic bead concentrator (Dynal), washed with 1X binding and washing (B&W) buffer (10 mM Tris-HCl, pH 7.5, 1 mM EDTA, 1 M NaCl) and TEN buffer (10 mM Tris-HCl, pH 7.5, 1 mM EDTA, 0.1 M NaCl), and incubated with 500 μg of cell nuclear extracts for 2 hrs at 4°C in the presence of 3 μg of poly (dI-dC). After washing with TEN buffer, bound proteins were eluted using 100 μl of 1X B&W buffer and analyzed by western blotting and mass spectrometry.

### Mice and tumor growth

C57BL/6J mice were obtained from Jackson Laboratory and maintained following the USDA guidelines. 6 to 8-week-old mice were injected with 5 × 10^5^ mEERL cells subcutaneously into the rear right flank (*n* = 10 per group). Tumor volume was measured twice a week and calculated using the equation: volume = (width^2^ X length)/2. Animals were euthanized when tumor volume reached 2000 mm^3^, as previously described (85). Conversely, mice were considered tumor-free when no measurable tumor was detected for 12 weeks. Survival graphs were calculated by standardizing for a tumor volume of 2000 mm^3^. The Michigan State University Institutional Animal Care and Use Committee (IACUC) approved experiments by the National Institutes of Health guidelines for using live animals.

## ACKNOWLEDGMENT

We thank members of the Pyeon laboratory for their valuable comments and suggestions. This work was supported in part by NIH R01 DE026125 and R01 DE029524 (D.P. and W.C.S.) and the Michigan State University Global Impact Initiative (D.P.). We thank Douglas Whitten from the Mass Spectrometry and Metabolomics Core at Michigan State University for providing mass spectrometry analyses.

Conceptualization, M.I.K. and D.P.; experimental design and methodology, M.I.K., C.Y., L.V., S.C., C.W., C.D.J., I.M.M., W.C.S., and D.P.; experiment, M.I.K., C.Y., L.V., S.C., and D.P.; writing original draft, M.I.K. and D.P.; review and editing, M.I.K., L.V., C.W., I.M.M., and D.P.

We declare no conflict of interest.

## FIGURE LEGENDS

**FIG. S1. The position and nucleotide sequence of the human *MARCHF8* promoter**. Shown are the *MARCHF8* promoter region (−90 to +160) with the positions of two E-boxes (red) and transcription start sequence (TSS, gray).

**FIG. S2. mRNA expression levels of FAS, TRAIL-R1, and TRAIL-R2 in HPV+ and HPV-cells**. The FAS, TRAIL-R1, and TRAIL-R2 mRNA expression levels in normal (N/Tert-1), HPV+ HNC (SCC-2, SCC-90, and SCC-152), and HPV-HNC (SCC-1, SCC-9, and SCC-19) cells (**A-C**) and N/Tert-1 cells expressing HPV16 E6, E7, or E6 and E7 (**D-F**) were quantified by RT-qPCR. The data shown are normalized by the GAPDH mRNA level as an internal control. All experiments were repeated at least three times, and the data shown are means ± SD. *P* values were determined by Student’s *t*-test. \**p* < 0.05, \*\**p* < 0.01, \*\*\**p* < 0.001.

**FIG. S3. Knockdown of *MARCHF8* expression increases FAS, TRAIL-R1, and TRAIL-R2 protein expression in HPV+ HNC cells**. HPV+ HNC (SCC2) cells were transduced with three lentiviral shRNAs against MARCHF8 (shR-MARCHF8 clones 3-5) along with scrambled shRNA (shR-scr). Protein expression of MARCHF8, FAS, TRAIL-R1, and TRAIL-R2 was determined by western blotting (**A**). Relative band density was quantified using NIH ImageJ (**B**). β-actin was used as an internal control. The data shown are means ± SD of three independent experiments. Cell surface expression of FAS (**C** and **D**), TRAIL-R1 (**E** and **F**), and TRAIL-R2 (**G** and **H**) proteins were analyzed by flow cytometry. Mean fluorescence intensities (MFI) of three independent experiments are shown (**D, F**, and **H**). *P* values were determined by Student’s *t*-test. \**p* < 0.05, \*\**p* < 0.01, \*\*\**p* < 0.001.

**FIG. S4. mRNA expression levels of FAS, TRAIL-R1, and TRAIL-R2 by knockdown of *MARCHF8* expression**. Two HPV+ HNC cell lines, SCC152 (**A-C**) and SCC2 (**D-F**), were transduced with five and three lentiviral shRNAs against MARCHF8 (shR-MARCHF8), respectively, or scrambled shRNA (shR-scr). The mRNA levels of FAS (**A** and **D**), TRAIL-R1 (**B** and **E**), and TRAIL-R2 (**C** and **F**) were quantified by RT-qPCR. The data shown are normalized by the GAPDH mRNA level as an internal control. All experiments were repeated at least three times, and the data shown are means ± SD. *P* values were determined by Student’s *t*-test. \*\*\**p* < 0.001.

**Table S1. MARCHF8 promoter binding proteins**

**Table S2. List of the antibodies**

**Table S3. List of the shRNAs**

**Table S4. List of the sgRNAs**

**Table S5. List of the oligonucleotides**

